# Image Downsampling Expedited Adaptive Least-squares (IDEAL) fitting improves intravoxel incoherent motion (IVIM) analysis in the human kidney

**DOI:** 10.1101/2022.06.09.495469

**Authors:** Julia Stabinska, Helge J. Zöllner, Thomas Thiel, Hans-Jörg Wittsack, Alexandra Ljimani

**Author notes:** Corresponding author: Julia Stabinska, Kennedy Krieger Institute, 707 N Broadway, Baltimore, MD 21205.

## Abstract

**Purpose:** To improve the reliability of intravoxel incoherent motion model (IVIM) parameter estimation for the diffusion-weighted imaging in the kidney using a novel Image Downsampling Expedited Adaptive Least-squares (IDEAL) approach.

**Methods:** The robustness of IDEAL was investigated using simulated diffusion-weighted MRI data corrupted with different levels of Rician noise. Subsequently, the performance of the proposed method was tested by fitting bi- and triexponential IVIM model to *in vivo* renal DWI data acquired on a clinical 3 Tesla MRI scanner and compared to conventional approaches *(Fixed D\** and *Segmented* fitting).

**Results:** The numerical simulations demonstrated that the IDEAL algorithm provides robust estimates of the IVIM parameters in the presence of noise as indicated by relatively low absolute percentage bias (sMdPB [%]) and normalized root-mean-square error (RMSE [%]). The analysis of the *in vivo* data showed that the IDEAL-based IVIM parameter maps were less noisy and more visually appealing than those obtained using the *Fixed D\** and *Segmented* methods. Further, the use of IDEAL for the triexponential IVIM modelling resulted in reduced cortical and medullary coefficients of variation (CVs) for all IVIM parameters when compared with *Fixed D*,* reflecting greater accuracy of this method.

**Conclusion:** The proposed fitting algorithm yields more robust IVIM parameter estimates and is less susceptible to poor SNR than the conventional fitting approaches. Thus, the IDEAL approach has the potential to improve the reliability of renal DW-MRI analysis for clinical applications.

## INTRODUCTION

Diffusion-weighted imaging (DWI) is a powerful MRI technique that measures microscopic random motion of water molecules in tissues and has been shown to provide unique information on renal microstructure and function^1^. Among the quantitative DWI-derived parameters, apparent diffusion coefficient (ADC) is the most widely used one and has been validated as a potential biomarker for different renal pathologies, e.g. renal artery stenosis^2^, kidney fibrosis^3,4^, renal lesions^5^, ureteral obstruction^6^, diabetic nephropathy^7^ and kidney graft rejection^8,9^. Nevertheless, the ADC value calculated from a mono-exponential model may not accurately reflect the self-diffusion behavior of water molecules in the renal tissue, as it is influenced by blood microcirculation and tubular flow. To separate the true tissue diffusion from pseudo-diffusion motion related to microperfusion, intravoxel incoherent motion (IVIM) model has been proposed^10^. Although the biexponential analysis of IVIM has been demonstrated to improve the interpretation of diffusion-attenuated MRI signal in the kidney compared to monoexponential ADC^11^, yet there is no uniform method of estimating the IVIM parameters. Furthermore, renal IVIM imaging is also challenged by limited data quality caused by the respiratory and cardiac motion and intrinsically low signal-to-noise ratio (SNR) of diffusion-weighted images, which may negatively affect the accuracy and reliability of the IVIM parameter estimates. Improved stability can be achieved by using segmented fitting approaches, which reduce the degree of freedom of the IVIM model. One commonly used method is the two-step fitting procedure in which the IVIM parameters are fitted stepwise after selecting a b-value threshold separating diffusion and pseudo-diffusion effects^12–14^. A challenging issue related to this type of fitting methods is the proper choice of the empirical b-value threshold above which the microcirculation-induced decay can be considered negligible. A simplified version of the segmented IVIM model fitting based on a low number of b-values (three or four) allows fast and numerically stable estimation of diffusion coefficients and perfusion fractions without, however, attempting to measure the pseudo-diffusion coefficients^15^.

Whereas previous studies have demonstrated the potential value of two-compartment IVIM-based DWI for assessing kidney pathologies^16–22^, more recently it has been hypothesized that the intra-tubular fluid compartment introduces a third component in the renal DW-signal^23–25^. Therefore, a rigid tri-exponential IVIM model has been proposed to distinguish between passive diffusion, tubular flow and vascular flow associated with the slow, intermediate, and fast (pseudo-) diffusion components, respectively. Nevertheless, the performance of the triexponential decay modelling is highly dependent on initial and boundary values used for the nonlinear data fitting. To overcome this limitation, either one or two pseudo-diffusion coefficients are often fixed. While this approach improves fit stability and sensitivity to physiological changes, it may also introduce bias to the perfusion fraction^24^. Therefore, there is a growing interest in developing novel fitting approaches to improve accuracy and repeatability of IVIM parameter estimation and facilitate clinical translation of IVIM-based imaging biomarkers.

In this study, we aimed at improving the reliability of rigid multi-exponential IVIM modelling of the human kidney by applying an Image Downsampling Expedited Adaptive Least-squares (IDEAL) fitting algorithm originally proposed for quantifying CEST MRI^26^. This approach utilizes combined spline interpolation and spatial downsampling to achieve higher SNR for iterative fitting. Moreover, it avoids arbitrary selection of initial values by tightly constraining the fit boundaries after initial fitting. We first tested the performance of the IDEAL fitting method on simulated kidney DWI data superimposed with different levels of noise. Subsequently, we applied the IDEAL fitting to quantify the IVIM parameters in healthy kidneys using bi- and triexponential IVIM models and compared the results with those obtained using the conventional fitting approaches, demonstrating the advantage of the IDEAL fitting for renal IVIM imaging.

## METHODS

### Multi-compartment IVIM models

The two-compartment IVIM model describes the diffusion-attenuated signal in the kidney as a two-exponential decay function expressed with the following formula:

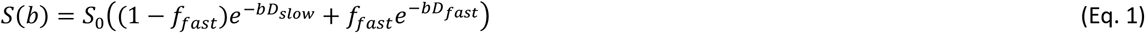

where *S*(*b*) is the signal intensity as a function of *b* (b-value), *S*_0_ it the signal in the absence of diffusion gradient sensitization, *D_slow_* is the diffusion coefficient of restricted water motion in renal tissue, *D_fast_* is the pseudo-diffusion coefficient of flow-based motion in tubules and blood capillaries, and *f_fast_* is the pseudodiffusion signal fraction.

As mentioned above, the three-compartment IVIM model may be preferable when interpreting renal DWI data as it enables the separation of two fast pseudodiffusion components, thus providing a more accurate assessment of both tubular flow and vascular perfusion in addition to renal tissue diffusion. If three (pseudo-) diffusion compartments are assumed, signal attenuation as a function of b-values is expressed by a tri-exponential equation:

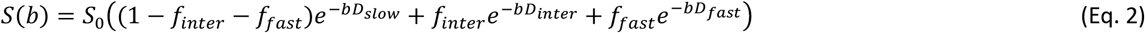

where *D_inter_* is the pseudo-diffusion coefficient of the intra-tubular fluid component, and *f_inter_* is the signal fraction of the tubular component, and *D_fast_* is the pseudo-diffusion coefficient of the blood component with the signal fraction *f_fast_*.

### Healthy subjects and patients

The study was approved by the local ethics committee, and written informed consent was obtained from all volunteers and patients (Ethics Committee of the Medical Faculty of Heinrich Heine University, Düsseldorf, Germany, study number: 2020-870). Nine healthy volunteers (5 females and 4 males; age range: 23.2 – 43.2 years; mean age: 30.4 ± 5.6 years) with no history of renal diseases were included in this study. Subjects were not given any restrictions regarding fluid or food intake.

### MRI

All MRI experiments were performed on a 3T MRI system (MAGNETOM Prisma, Siemens Healthcare, Erlangen, Germany) using an 18-channel torso array coil and a 32-channel spine coil. Anatomical T_2_-weighted images were acquired with a half-Fourier single-shot turbo spin echo sequence in the axial, coronal, and sagittal plane for planning the IVIM sequence. Diffusion-weighted images were obtained using a prototypical single-shot echo-planar imaging (EPI) sequence with a spectral-attenuated inversion-recovery (SPAIR) fat saturation and following parameters: TE/TR = 71/1900 ms; FOV: 370 x 370 mm^2^; acquisition matrix: 176 x 176; voxel size: 2.1 x 2.1 x 5.0 mm^3^; bandwidth: 1894 Hz/Px, GRAPPA factor: 2; 3 slices; 3 diffusion directions (3D diagonal); b-values (averages): (0 (3), 10 (3), 20 (3), 30 (3), 50 (3), 70 (4), 100 (4), 150 (4), 200 (4), 250 (4), 300 (4), 350 (4), 450 (5), 550 (5), 650 (5), 750 (5)) s/mm^2^. The DWI data acquisition was respiratory-triggered at the exhale phase of the respiratory cycle. The DWI scan duration was 4-6 minutes, depending on the subject’s respiration rate.

### Numerical phantom

A numerical phantom consisting of three kidney regions (renal cortex, renal medulla, and renal pelvis) and a background region was created in MATLAB (R2020a, Mathworks, Natick, MA) to test the robustness of the IDEAL fitting to noise. Parameters values for each of the kidney compartments were taken from the typical ranges reported in the literature and our experimental data^23^ and are given in Table 1. These values were kept constant in each kidney region, and the mean values of the parameters were calculated to define the background. The matrix size and b-values were the same as those used for *in vivo* experiments. The simulated data were corrupted with sixty-one different levels of Rician noise (standard deviation of the Rician noise distribution linearly sampled between 2 and 9.5) to yield datasets with SNRs between 91 and 19. SNR was determined from the b_0_ image as the ratio of the mean signal in parenchymal ROI to standard deviation of the noise.

**Table 1:** Summary of parameter values for each representative kidney compartment used for generating a numerical phantom.

| Compartment | Fraction (f) | | | (Pseudo-)Diffusion coefficient [$\times 10^{-3} \text{ mm}^2/\text{s}$] | | | $S_0$ |
| --- | --- | --- | --- | --- | --- | --- | --- |
| | $f_{\text{slow}}$ | $f_{\text{interm}}$ | $f_{\text{fast}}$ | $D_{\text{slow}}$ | $D_{\text{interm}}$ | $D_{\text{fast}}$ | |
| Cortex | 0.60 | 0.30 | 0.10 | 1.5 | 5.00 | 90 | 180 |
| Medulla | 0.70 | 0.22 | 0.08 | 1.6 | 5.50 | 110 | 160 |
| Pelvis | 0.30 | 0.40 | 0.30 | 1.1 | 9.00 | 300 | 215 |

### In vivo data analysis

Image registration based on a normalized mutual information similarity measure was applied to correct for spatial misalignment of DW-images of the volunteers using an in-house developed software based on ANTs (http://stnava.github.io/ANTs/). Cortical and medullary regions of interest (ROIs) in the right kidney were segmented manually by an experienced abdominal radiologist (AL, 9 years of experience) in ITK-SNAP (version 3.8.0)^27^. Subsequent data analysis was performed using custom-written MATLAB routines. The measured DW signal decays were fit to multiexponential functions with a constrained trust-region algorithm (MATLAB function *fit)* using different approaches: (i) an optimized tri-exponential fitting with fixed pseudo-diffusion coefficients (*Fixed D\**)^23^, (ii) a segmented bi-exponential fitting (*Segmented*)^28^ and (iii) an iterative IDEAL bi-exponential and tri-exponential fit *(IDEAL).* The source code for the IDEAL fitting is freely available on GitHub (https://github.com/stabinska/IDEAL). For the *Fixed D\** method, the three-compartment model were first fit to the ROI-based signals from the whole kidney volume to estimate D_fast_ and D_inter_ values, which were then averaged over all kidneys in all subjects. Next, the voxelwise fit for S_0_ and D_slow_, f_slow_, f_fast_, and f_inter_ with the fixed average pseudo-diffusion coefficients was performed. For the bi-exponential *Segmented* fitting, the threshold b-value of 200 s/mm^2^ was selected^29^. The initial values and boundaries for the signal fractions and (pseudo-) coefficients used for fitting are summarized in Table 2. In the *IDEAL* fitting, bi- and tri-exponential fitting of the averaged DW-signal from the images downsampled to 1 x 1, with the same loosely constrained lower and upper bounds as the conventional methods described above was performed to obtain initial values for a subsequent fitting step. These initial values and boundaries tightly constrained to 20% of the initial values for amplitude, 10% for the corresponding diffusion coefficients and 50% for the S_0_ image intensities were then used for voxel-wise fitting of the DW images downsampled to 2 x 2. The same steps were repeated for the DW images iteratively downsampled to 1 x 1, 4 x 4, 8 x 8, 16 x 16, 32 x 32, 64 x 64, 96 x 96, 128 x 128, 152 x 152 till the original acquisition matrix size of 176×176. The initial parameters of each voxel were determined by spatially interpolating the fitted parameters of the previous downsampled images using bilinear interpolation. The flow chart of the adapted *IDEAL* fitting procedure is displayed in Figure 1. To compare the performance of the triexponential IVIM modelling when using the *Fixed D\** and *IDEAL* algorithms, a representative dataset was corrupted with four different Rician noise levels with *σ* from 0 to 12. The highest noise level was chosen such that SNR of the DW-image at b value of 750 s/mm^2^ was higher than 5. The SNR was calculated as the ratio of the mean signal in the parenchymal ROI to the standard deviation of the noise in a ROI outside of the body.

**Figure 1:**
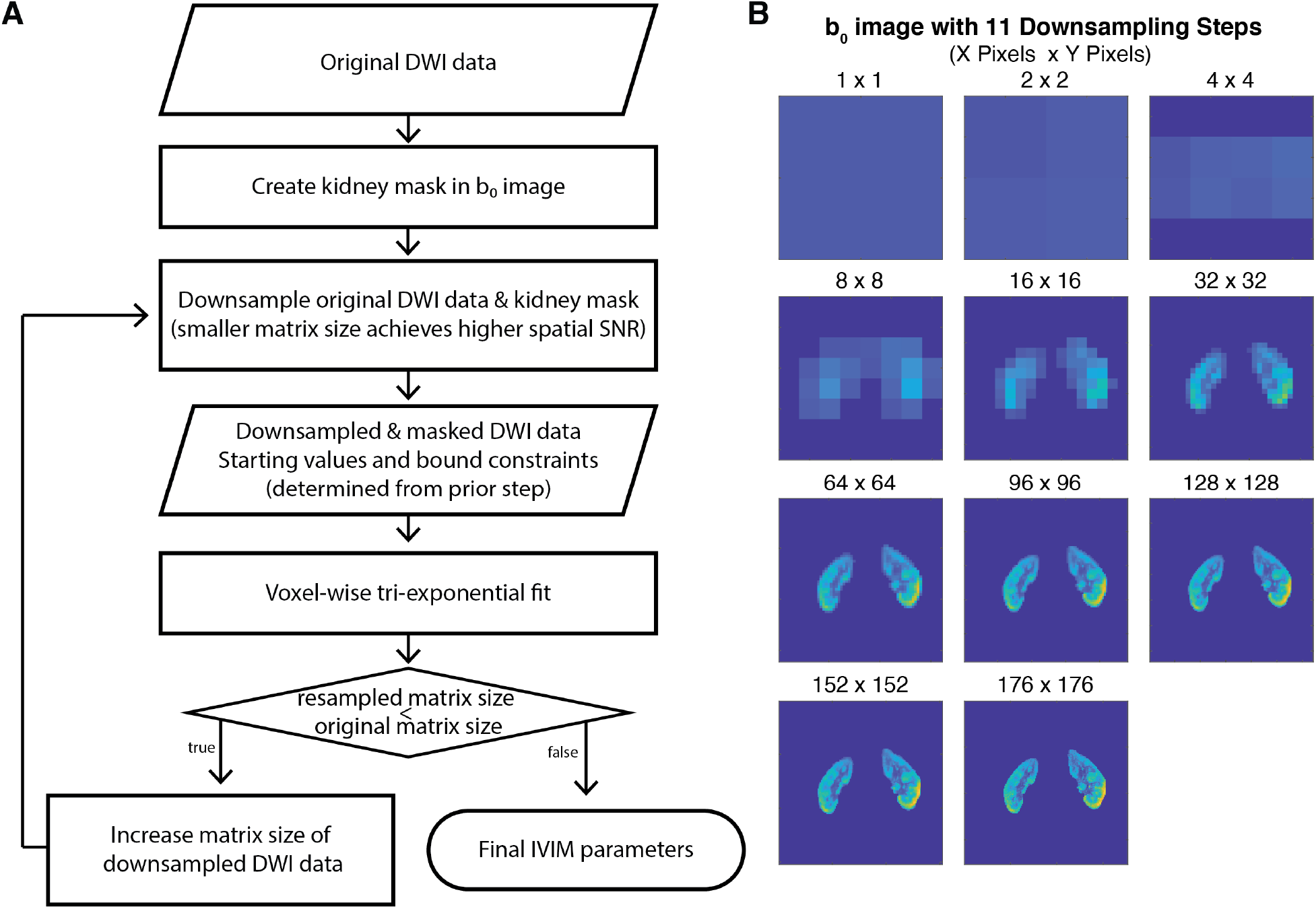
(A) Flow chart of data processing steps of the IDEAL fitting approach adapted for DWI. (B) Iteratively downsampled masked *in vivo* So images from a representative subject.

**Table 2:**
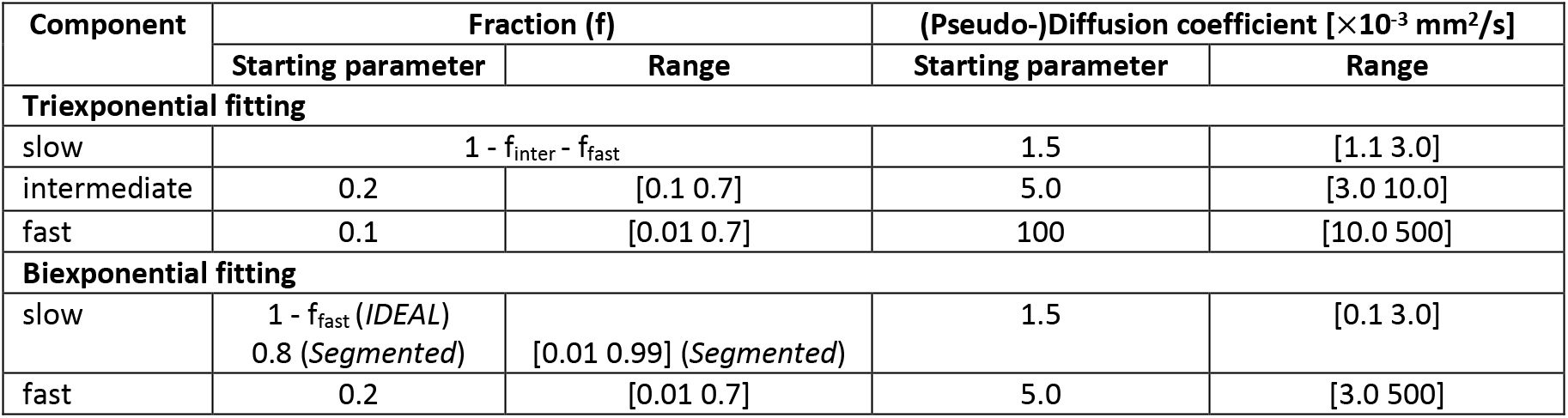
Summary of initial and boundary values used for tri- and biexponential IVIM fitting.

| Component | Fraction (f) | | (Pseudo-)Diffusion coefficient [$\times 10^{-3} \text{ mm}^2/\text{s}$] | |
| --- | --- | --- | --- | --- |
|  | Starting parameter | Range | Starting parameter | Range |
| **Triexponential fitting** |  |  |  |  |
| slow | $1 - f_{\text{inter}} - f_{\text{fast}}$ | | 1.5 | [1.1 3.0] |
| intermediate | 0.2 | [0.1 0.7] | 5.0 | [3.0 10.0] |
| fast | 0.1 | [0.01 0.7] | 100 | [10.0 500] |
| **Biexponential fitting** |  |  |  |  |
| slow | $1 - f_{\text{fast}}$ (*IDEAL*)<br>0.8 (*Segmented*) | [0.01 0.99] (*Segmented*) | 1.5 | [0.1 3.0] |
| fast | 0.2 | [0.01 0.7] | 5.0 | [3.0 500] |

### Statistical analysis

The quality of the IVIM parameter estimates obtained in the kidney phantom at different noise levels was verified by calculating the median percentage bias (sMdPB) defined as^30^:

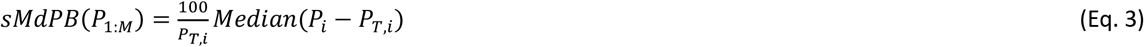

where *P_i_*, represents the IVIM parameter estimate in each pixel *i* (*i* = 1:M), while *P_T,i_* is the corresponding true value. Use of median was preferred over mean because it is less affected by outliers which in turn are biased by the choice of constraint boundaries^30^. The first and third quartiles were determined to provide a measure of variability and skewness of each IVIM parameter estimate. The systematic error of each estimate was measured as the normalized root-mean-square errors (RMSEs) between the fitted IVIM parameter and the ground truth.

Boxplots of cortical and medullary IVIM parameter estimates and coefficients-of-variation (CVs) obtained from the fit to the *in vivo* kidney data were generated for each fitting approach and evaluated quantitatively. The statistical analysis and visualization were performed using an open-source R-package “SpecVis” (version 1.0.0)^31^ available on Github (https://github.com/HJZollner/SpecVis) and R software (version 4.1.3, R Foundation for Statistical Computing, Vienna, Austria)^32^.

## RESULTS

### Numerical kidney phantom

Figure 2A displays the simulated parameter maps of the numerical kidney phantom with three different compartments (cortex, medulla and renal pelvis) and varying ground-truth values for each of these. The averaged sMdPB values for each parameter map obtained from the *IDEAL* fit at different noise levels are shown in Figure 2B. These results suggest that *IDEAL* provides robust estimates of the IVIM parameters in the presence of noise as indicated by low absolute percentage bias and relatively small IQR as represented by shaded areas. Reducing SNR to 45 resulted in a gradually increasing absolute bias for f_slow_, f_inter_, D_slow_ and D_inter_ with a maximum median value of about 20% for f_inter_ and −10% for f_slow_, D_slow_ and D_inter_ at SNR of 19. As expected, reduced SNR led to a gradual increase in normalized RMSE for all parameters with highest values for f_inter_, f_fast_ and D_fast_.

**Figure 2:**
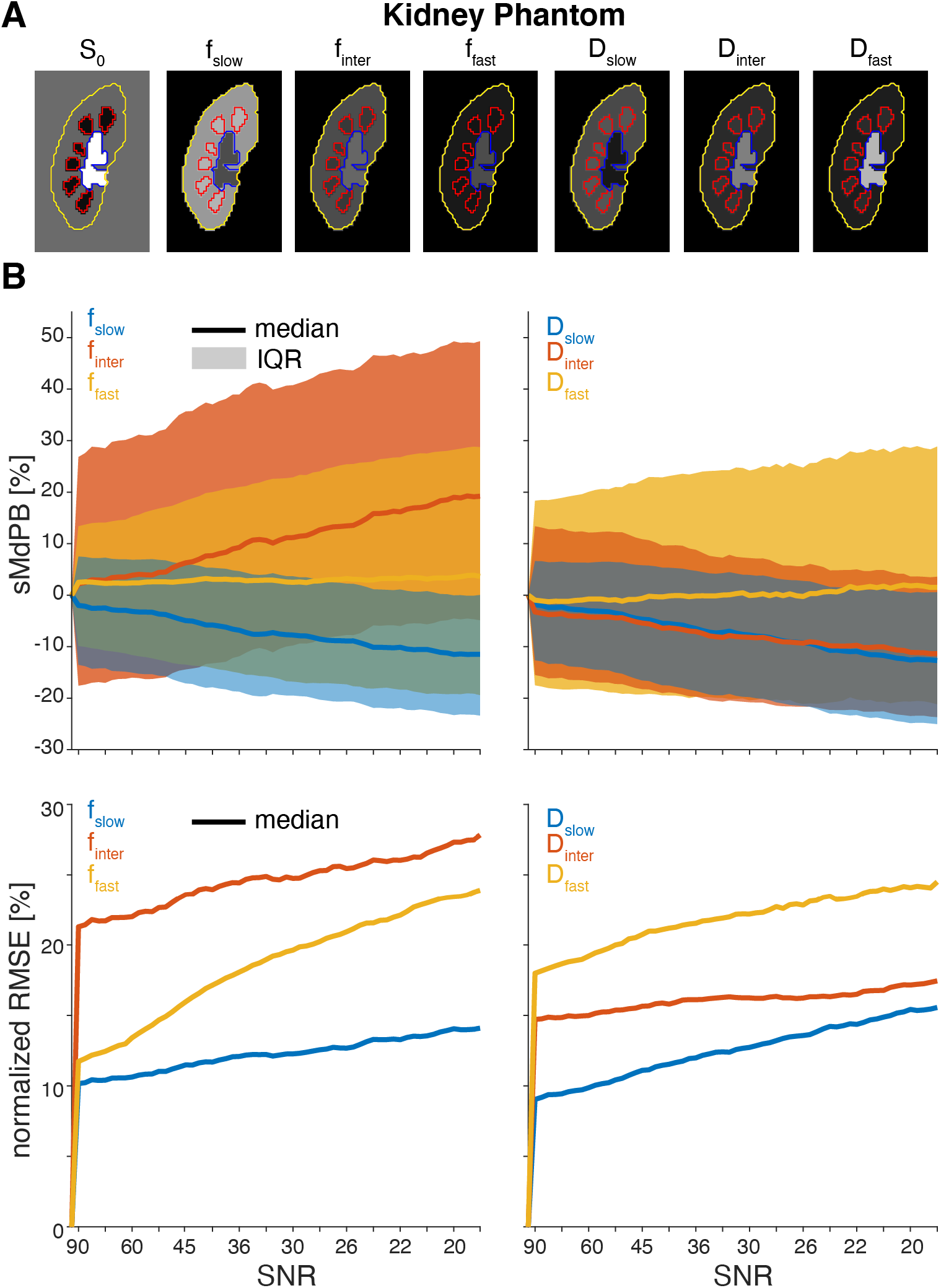
(A) S_0_ image and masked IVIM parameter maps of the simulated kidney phantom. Kidney, medulla, and pelvis masks are delineated yellow, red, and blues, respectively. (B) Median percentage bias (sMdPB) and normalized RMSE with respect to the ground-truth values for each parameter map at different SNR levels of the simulated data. The shown sMdPB and normalized RMSE values where smoothed using a moving average. The solid lines denote the median values and shaded areas represent the inter-quartile range. The SNR values are given for the S_0_ image.

### Triexponential IVIM analysis in the kidney

Exemplary signal fraction and diffusion coefficient maps obtained from the DWI data without (*σ* = 0) and with added noise (*σ* = 12) using the *Fixed D\** and *IDEAL* fitting approaches are displayed in Figure 3A and 3B. In general, the parameter maps fitted with *IDEAL* are less noisy than those calculated with *Fixed D*,* allowing better visualization of the complementary renal structures even at low SNR. Notably, the *IDEAL* method was superior at estimating D_slow_ than *Fixed D\** as shown in Figures 3C and 3D.

**Figure 3:**
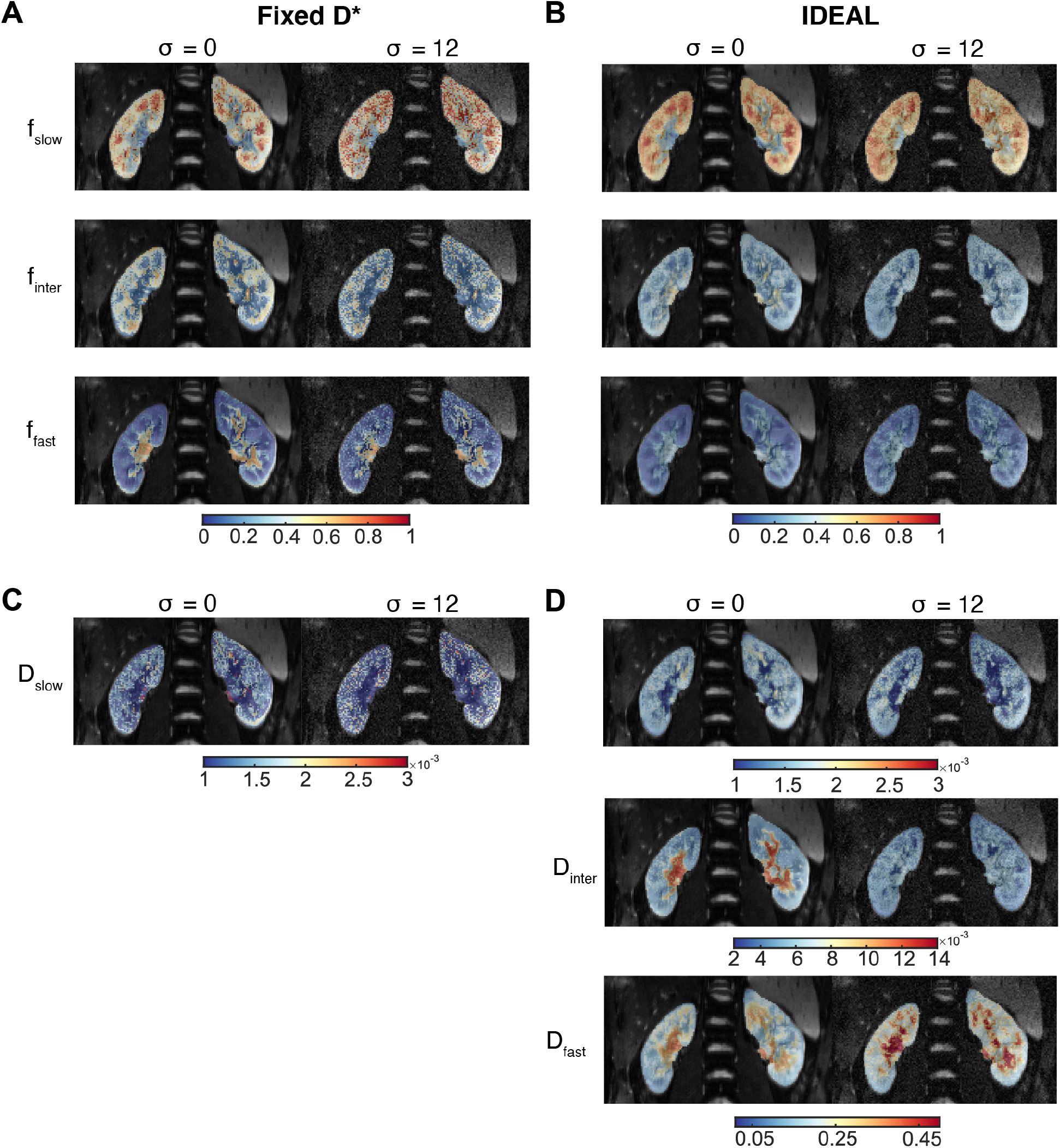
Signal fraction and diffusion coefficient maps obtained using triexponential IVIM modelling. (A, B) f_slow_, f_interm_, f_fast_ maps obtained with *Fixed D\** and *IDEAL* fitting from *in vivo* data without (σ = 0) and with added noise (σ = 12). (C, D) D_slow_ maps calculated with both fitting approaches, and (E, F) D_interm_ and D_fast_ fitted with IDEAL at two different noise levels.

Histograms in Figures 4A, 4B and 4C confirm these results, revealing overall narrower distributions of signal fraction and D_slow_ values calculated in the entire right kidney. The *Fixed D\** method resulted in a considerable number of voxels in which the estimated IVIM parameters reached or were close to the fit constraints. Further, the histograms in Figure 4D displaying the differences of the IVIM parameters obtained at different noise levels exhibit a more symmetrical distribution around zero and lower SDs (0.12 vs 0.19 for f_slow_, 0.11 vs 0.18 for f_iterm_, 0.04 vs 0.11 for f_fast_, and 2.39 x 10^-4^ mm^2^/s vs 4.60 x 10^-4^ mm^2^/s for D_slow_) with *IDEAL* compared to *Fixed D*,* indicating higher robustness of the former fitting approach to noise.

**Figure 4:**
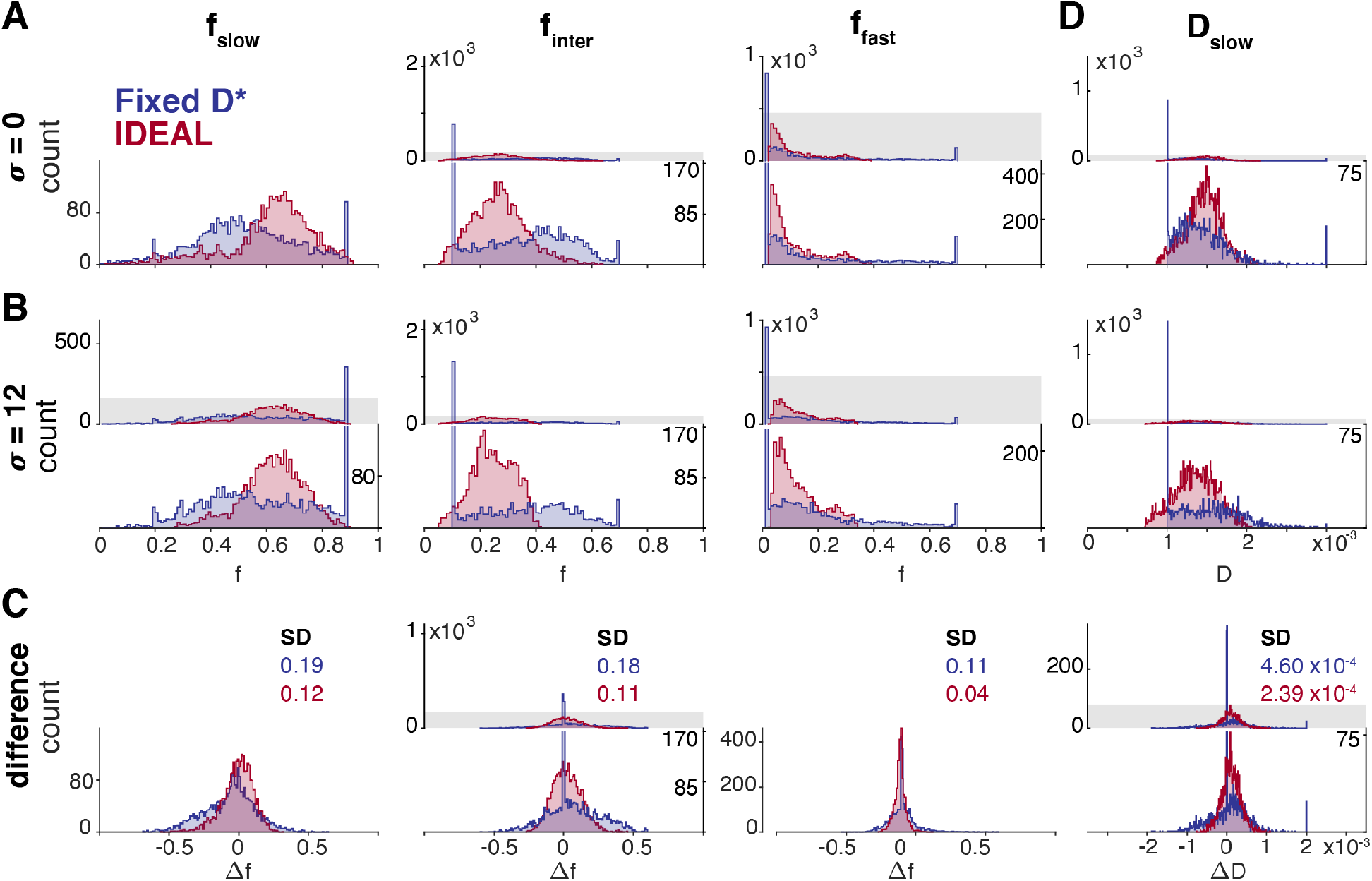
Histograms of IVIM parameter maps obtained using triexponential IVIM modelling with *Fixed D\** and *IDEAL* from *in vivo* data without and with added noise. (A) f_slow_, finter and f_fast_ maps at *σ* = 0 and (B) *σ* = 12, and (C) the corresponding D_slow_, maps. (D) Histograms of differences of the fitted IVIM parameter estimates for each voxel. For the panels with two axis the bottom axis shows magnification of the top axis (magnified region as gray shade).

Distribution of the median signal fractions and diffusion coefficient D_slow_ in the cortex and medulla of the right kidney obtained in all subjects are shown in Figure 5. There were no statistically significant differences in f_slow_, f_interm_, f_fast_ and D_slow_ values obtained from *Fixed D\** and *IDEAL* fitting in neither ROI (P > 0.158). At the same time, the cortical and medullary CVs of the fitted signal fractions and diffusion coefficient D_slow_ were significantly lower when *IDEAL* fitting was used (f_slow_: P = 0.013 and P = 0.009, f_interm_: P = 0.002 and P = 0.009, f_fast_: P = 0.0004 and P = 0.009, D_slow_: P = 0.009 and P = 0.0007 in the cortex and medulla, respectively). The IVIM parameters estimates fitted with *IDEAL* in the kidneys of all subjects reported as median and first and third interquartile [Q1, Q3] are summarized in Table 3.

**Figure 5:**
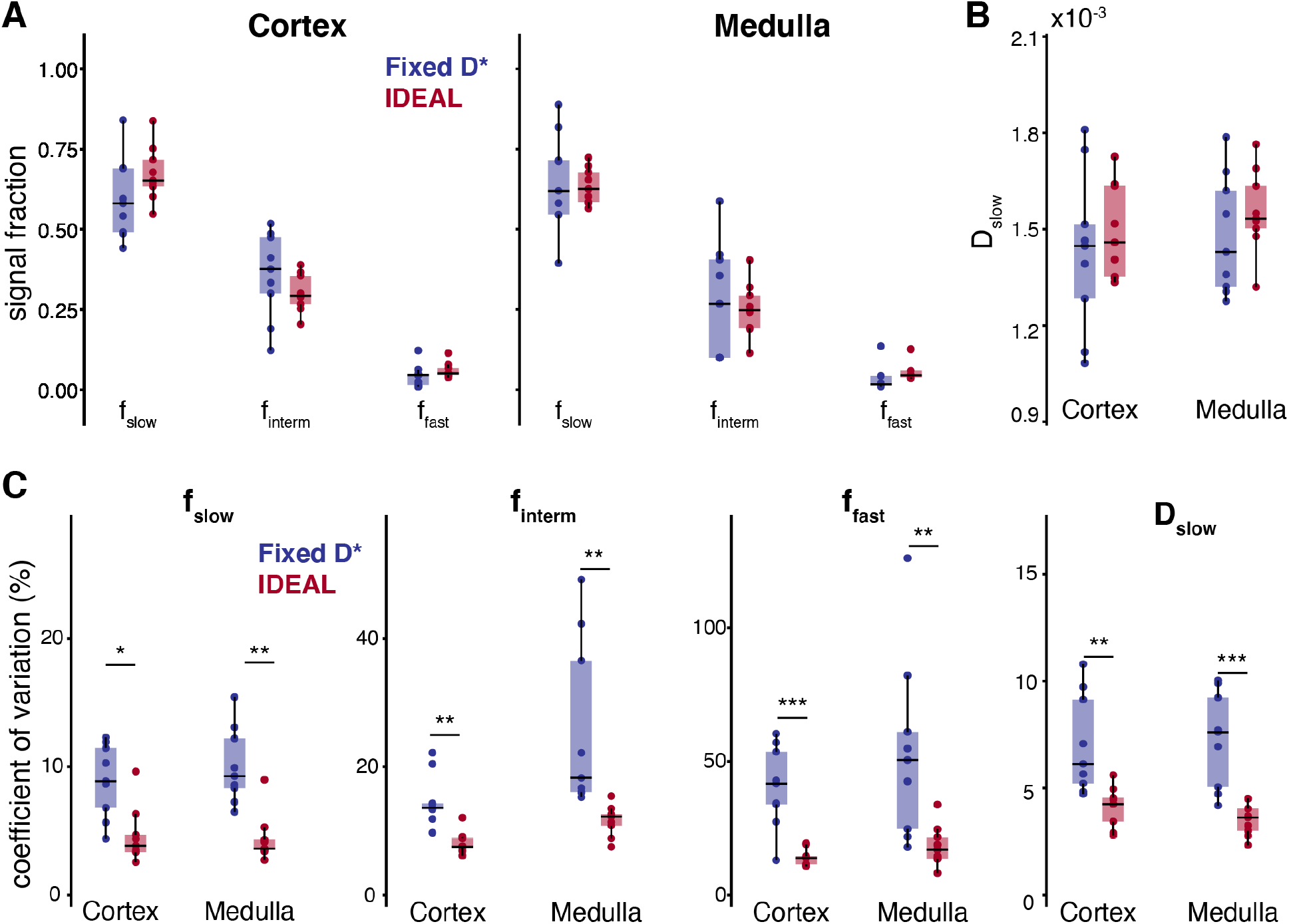
Distribution of median values of signal fractions obtained from triexponential IVIM modelling with *Fixed D\** and *IDEAL* fitting (A) in the cortex and medulla, and corresponding (B) D_slow_ values. (C) Distribution of coefficients of variation of the f_slow_, f_inter_, f_fast_ and D_slow_ parameters measured in the cortical and medullary ROIs in all subjects. Asterisks indicate significant differences with P < 0.05 = *, P < 0.01 = **, and P < 0.001 = ***.

**Table 3:** IVIM parameters fitted using the IDEAL algorithm. The values are presented as median, 25^th^ and 75^th^ percentile.

| Triexponential IVIM modelling |  |  |  |  |  |  |
| --- | --- | --- | --- | --- | --- | --- |
| | $f_{slow}$ | $f_{interm}$ | $f_{fast}$ | $D_{slow} [x10^{-3} \text{ mm}^2/\text{s}]$ | $D_{interm} [x10^{-3} \text{ mm}^2/\text{s}]$ | $D_{fast} [x10^{-3} \text{ mm}^2/\text{s}]$ |
| Cortex | 0.65 [0.63 0.72] | 0.29 [0.27 0.36] | 0.05 [0.05 0.07] | 1.46 [1.35 1.64] | 4.92 [3.93 5.96] | 87.37 [37.01 125.51] |
| Medulla | 0.63 [0.59 0.68] | 0.25 [0.19 0.29] | 0.05 [0.04 0.06] | 1.53 [1.50 1.63] | 5.24 [3.94 5.75] | 134.57 [47.70 165.49] |
| Biexponential IVIM modelling |  |  |  |  |  |  |
| | $f_{slow}$ | $f_{fast}$ | $D_{slow} [x10^{-3} \text{ mm}^2/\text{s}]$ | | $D_{fast} [x10^{-3} \text{ mm}^2/\text{s}]$ | |
| Cortex | 0.82 [0.81 0.86] | 0.18 [0.14 0.19] | 1.69 [1.63 1.90] |  | 10.85 [9.33 13.65] |  |
| Medulla | 0.85 [0.85 0.88] | 0.15 [0.12 0.15] | 1.79 [1.71 1.93] |  | 13.89 [9.45 26.78] |  |

### Biexponential IVIM analysis in the kidney

Exemplary IVIM parameter maps obtained from the biexponential fitting using the *Segmented* and *IDEAL* fitting approaches are presented in Figure 6. While the *Segmented* maps exhibit spatial noise, the *IDEAL* fit yields more visually appealing signal fraction and especially (pseudo-) diffusion coefficient maps. The boxplots displayed in Figure 7 show lower variance of the median f_slow_, f_inter_, f_fast_ and D_fast_ estimates in the cortex and medulla for *IDEAL* compared to the *Segmented* fitting. At the same time, there were no significant differences between the tested approaches in any of the IVIM parameters (P > 0.188). Further, the medullary CVs (calculated for each subject) for all IVIM parameters were significantly lower with the *IDEAL* approach, while the cortical CVs of the two considered fitting methods were not significantly different (P > 0.058).

**Figure 6:**
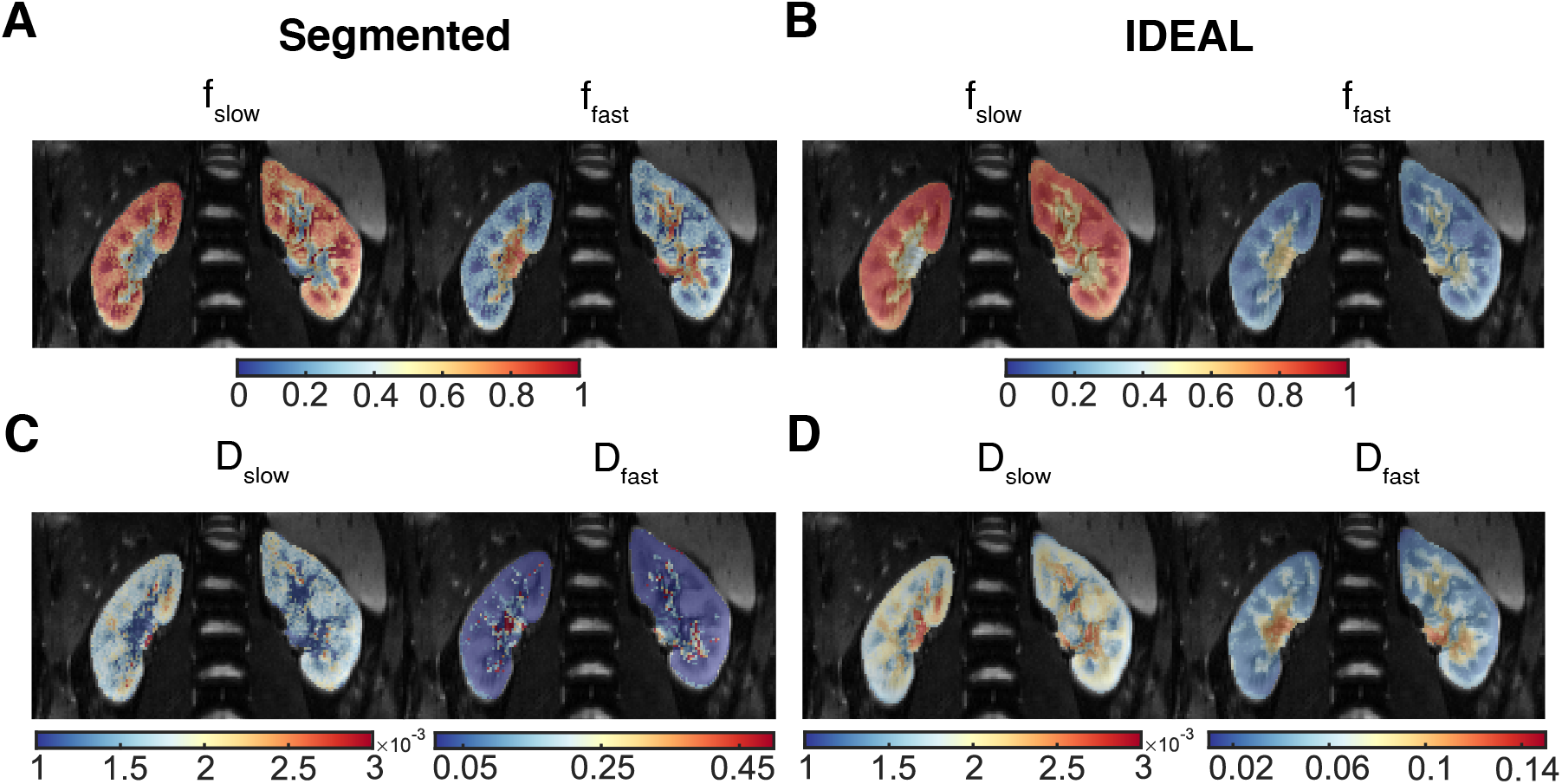
Signal fraction and diffusion coefficient maps obtained using biexponential IVIM modelling. (A, B) f_slow_ and f_fast_ maps obtained with *Segmented* and *IDEAL* fitting from *in vivo* data, and corresponding (C, D) D_slow_ and D_fast_ maps calculated using the two fitting approaches. Note the different scales of the D_fast_ maps for the *Segmented* and *IDEAL* methods.

**Figure 7:**
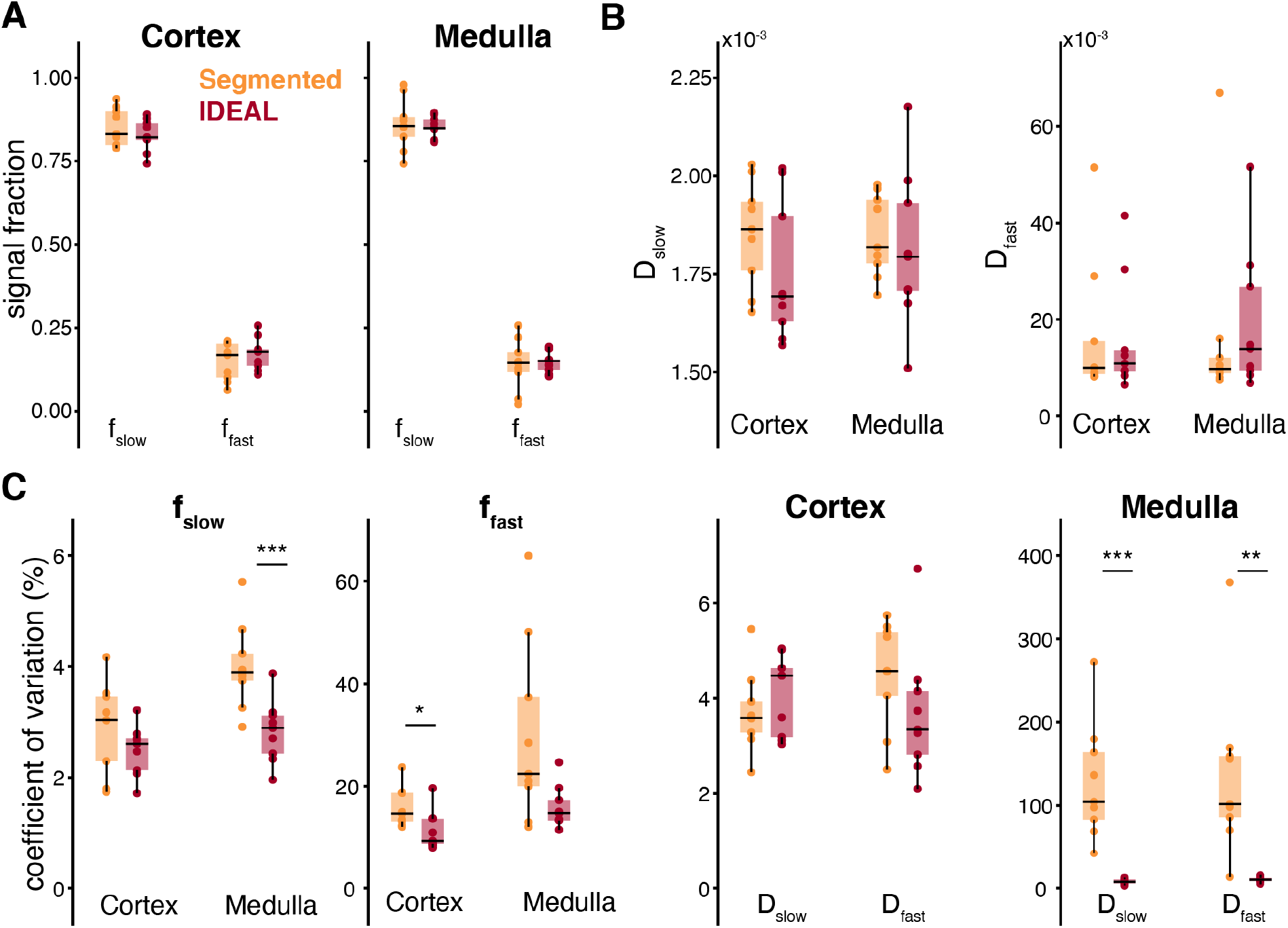
Distribution of median values of signal fractions obtained from biexponential IVIM modelling with *Segmented* and *IDEAL* fitting (A) in the cortex and medulla, and corresponding (B) D_slow_ and D_fast_ values. (C) Distribution of coefficients of variation of the f_slow_, f_fast_, D_slow_ and D_fast_ parameters measured in the cortical and medullary ROIs in all subjects. Asterisks indicate significant differences with P < 0.05 = *, P < 0.01 = **, and P < 0.001 = ***.

## DISCUSSION

In the present study, the IDEAL fitting algorithm proposed by Zhou et al.^26^ was adopted for DWI and compared to commonly used fitting approaches for multi-exponential IVIM modelling of the human kidney. The proposed novel approach utilizes high SNR of spatially downsampled images for iterative fitting with tightly constrained boundaries, which are updated after each iteration until the original image resolution is reached.

Numerical simulations in the kidney tissue-like phantom and analysis of the noise-corrupted *in vivo* DWI data revealed that the use of the IDEAL algorithm allows more reliable and more robust fitting of IVIM parameters, and is less susceptible to poor SNR than the conventional methods. The IVIM parameter maps generated using IDEAL were overall less noisy with fewer/no apparent outliers compared to those obtained using the *Fixed D\** approach. Notably, application of the IDEAL algorithm for triexponential IVIM analysis resulted in reduced CVs for all IVIM parameters in the cortex and medulla when compared with *Fixed D*,* reflecting greater accuracy of the proposed method. When applied for fitting the biexponential IVIM model, IDEAL yielded significantly lower CVs for medullary f_slow_, cortical f_fast_ and medullary D_slow_ and D_fast_. This superiority can be also observed in the parameter maps, especially D_slow_ and D_fast_ maps, which are less speckled and provide improved visualization of the complementary renal structures in comparison to the *Segmented* approach.

In the past years, many different approaches i.e., signal averaging and denoising techniques^33,34,34^ have been proposed to overcome the inherent limitation of IVIM imaging related to the comparatively low SNR of the DWI data especially at high b-values. In particular, a recent study of Kurugol et al. demonstrated that using a spatial homogeneity prior in a Markov Random Field framework improves the robustness of the parameter estimations of a probability distribution model of incoherent motion (SPIM)^35^. In fact, this and other works showed that accounting for spatial homogeneity not only increases SNR but also enforces appropriate boundaries between homogenous regions when these boundaries are present^36,37^. Note that the proposed IDEAL fitting for IVIM exploits the concept of spatial homogeneity by using spline interpolation for determining the initial values for IVIM fitting in each voxel.

The commonly used IVIM fitting approaches such as the segmented method or tri-exponential fitting with fixed pseudodiffusion coefficients improve the fit stability by reducing the degree of freedom of the IVIM model. Although both these methods generally outperform the conventional unconstrained voxel-wise fitting with Levenberg-Marquardt algorithm, previous study demonstrated that fixing D_fast_ (and D_interm_) decreases the precision of the IVIM model as the changes in the pseudodiffusion coefficients affect the signal fractions instead. Hence, less constrained yet robust novel fitting methods are needed to enable more accurate characterization of the (pseudo-) diffusion in the renal tissue.

A very promising class of algorithms that have been proposed to fit the IVIM model are Bayesian probability-based methods. These approaches regularize the fitting problem to mitigate overfitting, and assume spatial dependencies between voxels for improved fitting performance^38–41^. Despite their comparatively high accuracy and precision as well as low inter-subject variability, Bayesian methods are relatively time-consuming and may lead to biased estimates of the pseudo-diffusion coefficients^42^. Compared with these approaches, the proposed IDEAL algorithm enables vastly faster parameter estimation without the necessity of specifying a prior distribution. Lately, more advanced approaches based on e.g. convolutional neural networks (CNN)^42–46^ and homology-based topological analysis^47^ have shown some promise for solving the mathematically ill-posed problem of computing IVIM parameters derived from the biexponential model. While using NN for IVIM fitting is several orders quicker than using the Bayesian methods, the networks may require retraining for different acquisition protocols or pathologies. The use of the IDEAL fitting represents a relatively simple alternative to the NN-based methods as it does not require any training and recalibration. Additional studies are needed to directly compare the performance of these methods for renal/abdominal DWI MRI especially for fitting the tri-exponential IVIM model.

Consistent with the morphological and physiological characteristics of the kidney, evidence of the existence of an additional intermediate pseudo-diffusion component in the renal tissue has been provided in few recent studies in which less constrained, data-driven non-negative least squares (NNLS) continuum modelling has been utilized to determine the number of diffusion compartments present in the renal tissue^25,48^. Although the NNLS approach has the potential to reflect the complex nature of water diffusion in the kidney more accurately than the rigid multi-exponential models, it is also susceptible to low SNR leading to biased estimates and noisy parameter maps^49^.

This study has several limitations worth noting. Our cohort of nine subjects was small and no patients with renal pathologies were examined. In addition, due to lack of a reference method and/or histological assessment it was not possible to validate the DWI-derived parameters in any way and correlate these with histological findings. Moreover, food and water intake were not restricted, which might have led to a higher variability of diffusion parameters^50^. Further, moderate spatial resolution of DWI may have caused partial volume averaging between cortex and medulla. Finally, we only analyzed a single slice of the right kidney which does not necessarily provide representative information about all kidney (pseudo-)diffusion compartments.

## CONCLUSION

In summary, the present study demonstrates the utility of IDEAL fitting algorithm for IVIM modelling of the kidney. In fact, the IVIM parameter estimates obtained using this novel approach are in good overall agreement with recent literature^23,29^. We showed that renal IVIM modelling using the IDEAL algorithm yields more robust IVIM parameter estimates as compared to conventional fixed and segmented fitting approaches. Further studies are warranted to assess the performance of this novel technique in differentiating between normal and pathological renal tissue or when detecting changes under different physiological conditions within the context of longitudinal studies.

## Acknowledgement

The study was supported by a grant from the German Research Foundation (DFG)(Project number: 408765040).

